# Ocular following eye movements in marmosets follow complex motion trajectories

**DOI:** 10.1101/2023.03.11.532200

**Authors:** Jagruti J. Pattadkal, Carrie Barr, Nicholas J. Priebe

## Abstract

Ocular following eye movements help stabilize images on the retina and offer a window to study motion interpretation by visual circuits. We use these ocular following eye movements to study motion integration behavior in the marmosets. We characterize ocular following responses in the marmosets using different moving stimuli such as dot patterns, gratings, and plaids. The marmosets can accurately track motion along different directions and exhibit spatial frequency and speed sensitivity that closely matches the sensitivity reported in neurons from their motion selective area MT. Marmosets are also able to track the integrated motion of plaids, with tracking direction consistent with intersection of constraints model of motion integration. Marmoset ocular following responses are similar to responses in macaques and humans with certain species-specific differences in peak sensitivities. Such motion sensitive eye movement behavior in combination with direct access to cortical circuitry makes the marmoset model well suited to study the neural basis of motion integration.

**Significance statement:** Ocular following is a reflexive eye tracking behavior in response to large visual field motion. It reflects the properties of underlying motion sensing circuits. One of the primary motion sensing areas in primates is area MT. In the primate species of marmosets, this and other cortical areas are easily accessible due to their lissencephalic brain. We demonstrate ocular following behavior in the marmosets for simple and complex motion trajectories and describe its characteristics. We then use ocular following to distinguish between different motion integration models. Our results show the utility of ocular following to study the neural basis for motion sensing in marmosets.

## Introduction

A critical function of sensory processing is to integrate distinct information from our environment to generate a global representation to guide our behavior. As information flows through the visual system, transformations occur that integrate multiple signals to constrain the interpretation of our visual environment. An example of this process occurs in the visual system when inferring the motion of complex objects. This inference requires the integration of multiple visual motion signals, each providing a constraint on the interpretation of the global object motion; seen for instance in our interpretation of overlapping moving grating stimuli – a plaid that appears to move in the direction distinct from either of the grating stimuli. The emergent representation of this pattern motion provides us with a model system to examine the neural circuitry that underlies signal integration in cortex.

Motion signal integration can occur following different principles such as vector averaging (VA) or intersection of constraints (IOC, see below). Motion of the individual grating components in a plaid stimulus is ambiguous and could correspond to set of different motion interpretations. This ambiguity is known as the ‘aperture problem’. For a single grating, the interpretations of its motion lie along a line in the velocity space and we generally perceive the motion corresponding to the slowest velocity. Overlapping gratings moving in different directions constrain our interpretation of motion: in this case, plaid motion can be computed based on the vector average (VA) of the motion direction of the components (Yo & Wilson, 1992), or by considering a more complex model in which each motion component contributes a constraint line, and the intersection of constraints lines from multiple components yields our percept (intersection of constraints – IOC (Adelson & Movshon, 1982)). For cases in which the two gratings are moving at the same speed (“Type I Plaids”), the VA and IOC models yield the same direction. Plaid stimulus where two motion components move at different speeds (“Type II Plaids”), can yield IOC predictions that differ from VA predictions by lying outside the two motion components. The VA predictions always lie between the motion components. Humans generally perceive motion in the IOC direction, though this percept depends on the contrast of the motion components, and duration of stimulus presentation.

We can infer the interpreted motion by using the eye movements evoked in response to the moving patterns. Brief presentations of large moving stimuli evoke reflexive ocular following eye movements that aid to stabilize the visual world on the retina (Kawano & Miles, 1986; Miles, 1998; Miles et al., 1986; Miles & Kawano, 1986). Ocular following in macaques depends on activity in motion selective neurons in the dorsal visual pathway including areas like MT and MST (Kawano et al., 1994; Takemura et al., 2007). The direction of human ocular following eye movements in response to the moving plaids is biased toward the IOC direction, particularly at long latencies (Masson & Castet, 2002). The eye movement behavior can be used to gain insights into the motion integration computation performed by the motion pathway. We examine ocular following responses in the primate model of marmosets. These animals are particularly amenable to studying motion computations due to their lissencephalic brain. The motion selective areas like MT and MST all lie on the brain surface and are easy to access for neural recordings (Pattadkal et al., 2022), unlike in the macaque brain. We demonstrate that marmosets are a good model for studying ocular following behavior and describe the ocular following response properties in these animals using different stimuli such a field of moving dots, gratings as well as type II plaids. Marmoset ocular following responses match those found in humans, though the spatial parameters eliciting peak responses differ in ways that reflect the spatial resolution of the marmoset visual system.

## Methods

Eye movements were collected from four marmoset subjects (3 males, 1 female) who were trained to fixate small spots or faces. Prior to training a head post was affixed to the skull in a sterile procedure performed in isoflurane anesthesia. The titanium head post was affixed to the skull using Metabond (Parkell, Edgewood, New York). Once the marmosets recovered from the head post procedure they were trained to receive juice reward for fixating small spots or faces (Mitchell et al., 2015). Subjects were maintained on food control to provide motivation in behavioral tasks with their weight ranging from 5-10% of baseline. All procedures conformed to NIH guidelines and the guidelines of the [Author University] Institutional Animal Care and Use Committee.

### Experimental procedures

Eye movements were tracked using the Eyelink 1000 plus system for IR eye tracking. The data was collected from the right eye in pupil-CR mode at 1kHz. Stimuli were displayed on a gray background screen (mean luminance 127). The task control and data collection were done using Maestro software (https://sites.google.com/a/srscicomp.com/maestro/). Animals were initially required to fixate on a target 5 degrees from the central fixation for 500-800 ms and then the target was shifted to the center of the screen. Animal was required to fixate on center target for 250-300 ms with a 220 ms grace period. Once the animal made this saccade to the central target, the target was extinguished and a dot field, grating or plaid stimulus presented and moved for 200-400 ms. No requirement for eye movement was made for the motion stimulus and if the animal performed the initial saccadic eye movement from the distal to central target, they received a juice reward.

All stimuli were presented on a FlexScan T761 50 cm (19 inch) Class Color Display monitor with a refresh rate of 85 Hz. This monitor was placed 50 cm away from the marmoset and field of view was 35 by 28 degrees.

### Preprocessing of eye movements

We observed that baseline eye positions following fixation to second target tended to drift. This may be similar to the glissades following saccades reported in previous studies (Kawano & Miles, 1986). Averaging the responses to the saccades only, without a motion stimulus (blank), can be used to cancel these drifts. We however observed that this drift was not common across all trials, even when the saccade direction was the same. The amount of drift in the initial period, before the moving target can affect eye movement, is not related to the motion response of the eye movement later in the trial. But we have chosen to exclude trials which show this drift, by enforcing a threshold criterion for position deviation. We smoothed horizontal and vertical eye positions using a median filter with an order of 30. The maximum deviation is calculated from these smoothed positions during an interval starting 55 ms before motion onset to 25 ms after motion onset. Any trial with position deviation exceeding 0.2 degrees in either vertical or horizontal position was excluded from our data. In addition, trials with any saccade onset in a period of 150 ms after motion onset were also excluded from our analysis.

### Eye velocity calculation

To compute the eye velocity, eye position trace collected at 1kHz was median filtered with an order of 25. The trace was then differentiated at 50 and 25 Hz and the mean was used as eye velocity estimate (Mitchell et al., 2015).

### Latency calculation

In order to estimate the latency of ocular following responses, we first obtained an average eye velocity trace. This average was calculated across trials, sampled with replacement. The total number of trials averaged equals the number of trials in the dataset for that condition. The average eye velocity *e_v_* is then fitted with a threshold linear function of the form:

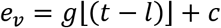

Where *g* is a gain factor, *t* is time of the trace being 0 at motion onset, *l* is the latency of the velocity trace and *c* is the offset. We bootstrap this latency measurement, by repeating the process 1000 times and use the median of the generated distribution as the mean latency for that condition. Latencies used for each test were calculated either for the conditions evoking strongest responses or averaged across different conditions when the responses had similar magnitude.

### Open loop interval and response amplitude calculation

We have restricted our ocular following response amplitude measurements to the open loop period from motion onset. Open loop interval was estimated to end at twice the latency estimate (Masson & Castet, 2002). Amplitude of ocular following response was calculated by subtracting the average velocity before the start of the response (latency) from the average velocity at the end of the open loop interval. The average velocities were computed in 15 ms period before the latency and before the end of the open loop interval respectively.

### Sinusoidal fit for dependence of OFR on motion direction

The horizontal and vertical eye velocities were fitted with sinusoidal function:

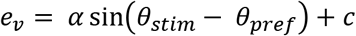

Where *e_v_* is the average eye velocity, *α* is the amplitude, *θ_stim_* is the stimulus direction, *θ_pref_* is the preferred stimulus direction and *c* is the offset term.

Circular correlation between ocular following angle (*θ_em_*) and stimulus direction (*θ_stim_*) was computed using the following formula (Batschelet, 1981):

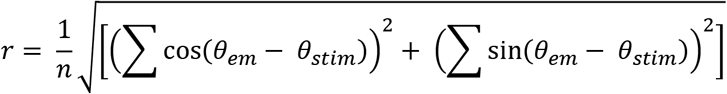

Where n is the number of conditions.

### Gaussian fit for dependence on speed or spatial frequency

The OFR amplitude dependence on speed or spatial frequency was fit with a Gaussian function:

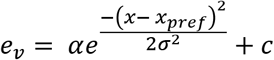

Where *e_v_* is the average eye velocity, *α* is the gain factor, *x* is the stimulus speed or spatial frequency, *x_pref_* is the preferred stimulus speed or spatial frequency, *σ* is the standard deviation of the Gaussian and *c* is the offset term. The fits were calculated on a log scale.

## Results

We aim to characterize ocular following responses (OFRs) and determine how ocular following responses follow complex motion trajectories in the marmosets. Ocular following responses (OFRs) have been characterized in a number of mammals, including rodents, macaques and humans but the visual parameters required to evoke these smooth eye movements differ across species. Because these eye movements have not previously been characterized in marmosets, we initially describe the visual parameters that elicit OFRs.

### Ocular following responses to simple motion stimuli

To evaluate the visual parameters that evoke OFRs we used a saccade-initiated paradigm in which animals are required to make a saccadic eye movement from a 5 degree eccentric target to a central target, after which a motion pattern was presented (Fig. 1A). We used this saccade initiated paradigm because it reduced the saccade instances within few hundred ms after motion onset. This is the interval used for analyzing ocular following responses and trials with saccades during this interval were discarded (see methods).

**Figure 1:**
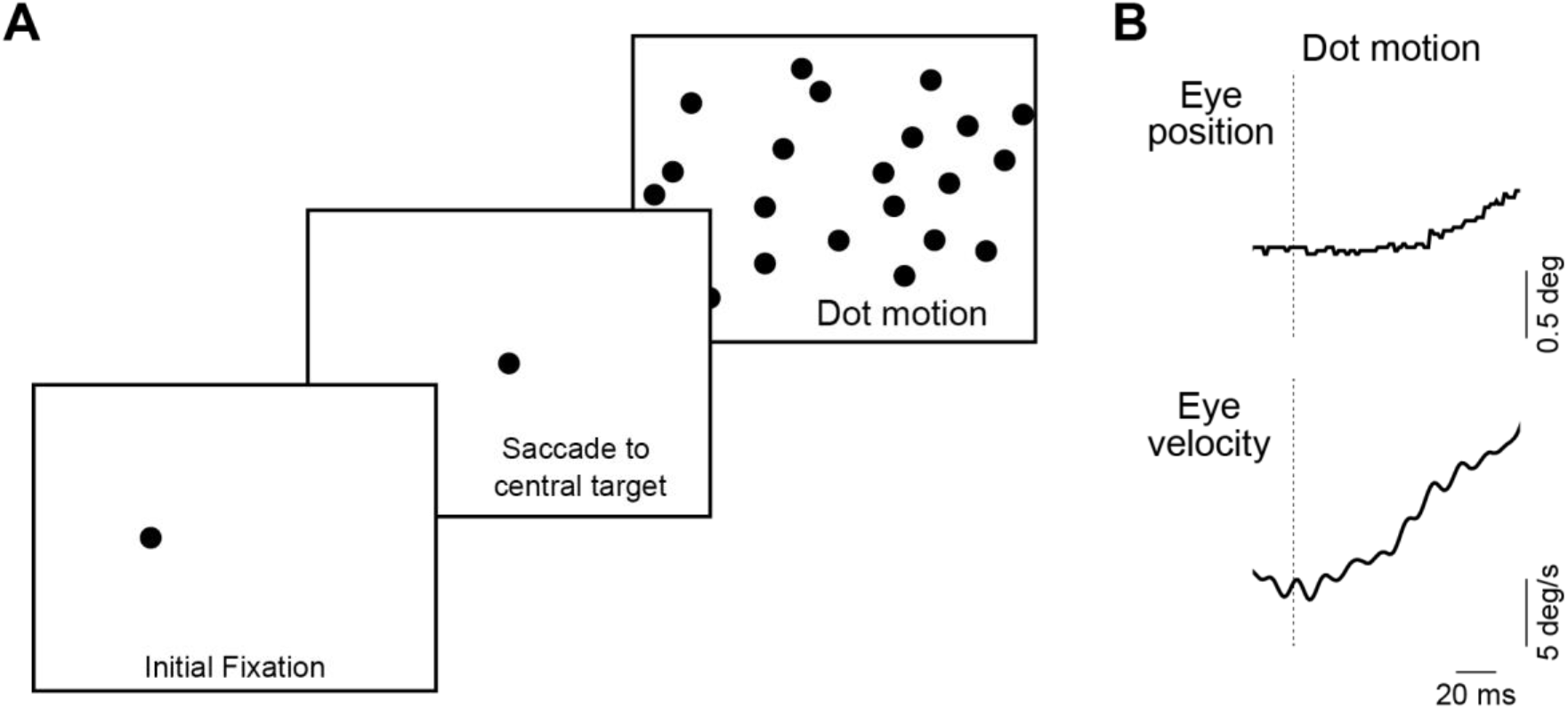
Ocular following task and example eye movement. A) Schematic of the ocular following task. Trials began with an initial fixation target, 5 degree away from the center of the screen. Animals were then required to make a saccade to a central fixation target. Following this fixation, motion stimulus turned on, which was either a field of dots, a grating, or a plaid. B) Example ocular following response in a marmoset for a dot field motion. Upper panel shows the eye position trace and lower panel shows the eye velocity trace. Dashed vertical line indicates the start of motion stimulus.

We initially presented moving patterns that were composed of randomly placed dots which varied in size from 0.6 to 1.6 degrees. In response to rightward motion of the dot pattern at 25 deg/s, clear smooth movements are evoked following the saccade to the central target (Fig. 1B). Smooth eye movements were initiated rapidly after the visual pattern movement (median latency animal A: 57 ms, median latency animal B: 64 ms). To quantify the mean OFRs, we measured the mean evoked eye velocity in the open-loop interval from a 25 to 40 ms period post latency. For rightward motion, the mean OFR was 4.5 +/- 0.2 s.e.m. deg/s in animal A and 3 +/- 0.4 s.e.m. deg/s in animal B, a comparable amplitude to those observed in the macaque. OFR gain was comparable for both animals (mean gain 0.18 +/- 0.008 in animal A, mean gain 0.12 +/- 0.016 in animal B).

Changing the direction of dot motion systematically shifted the horizontal and vertical eye velocity components (Fig. 2A), and no clear biases were observed in either marmoset. The mean eye tracking direction of the ocular following responses closely matched the presented the stimulus motion direction, for both animals (Fig. 2B, circular correlation 0.99 for animal 1, 0.96 for animal 2).

**Figure 2:**
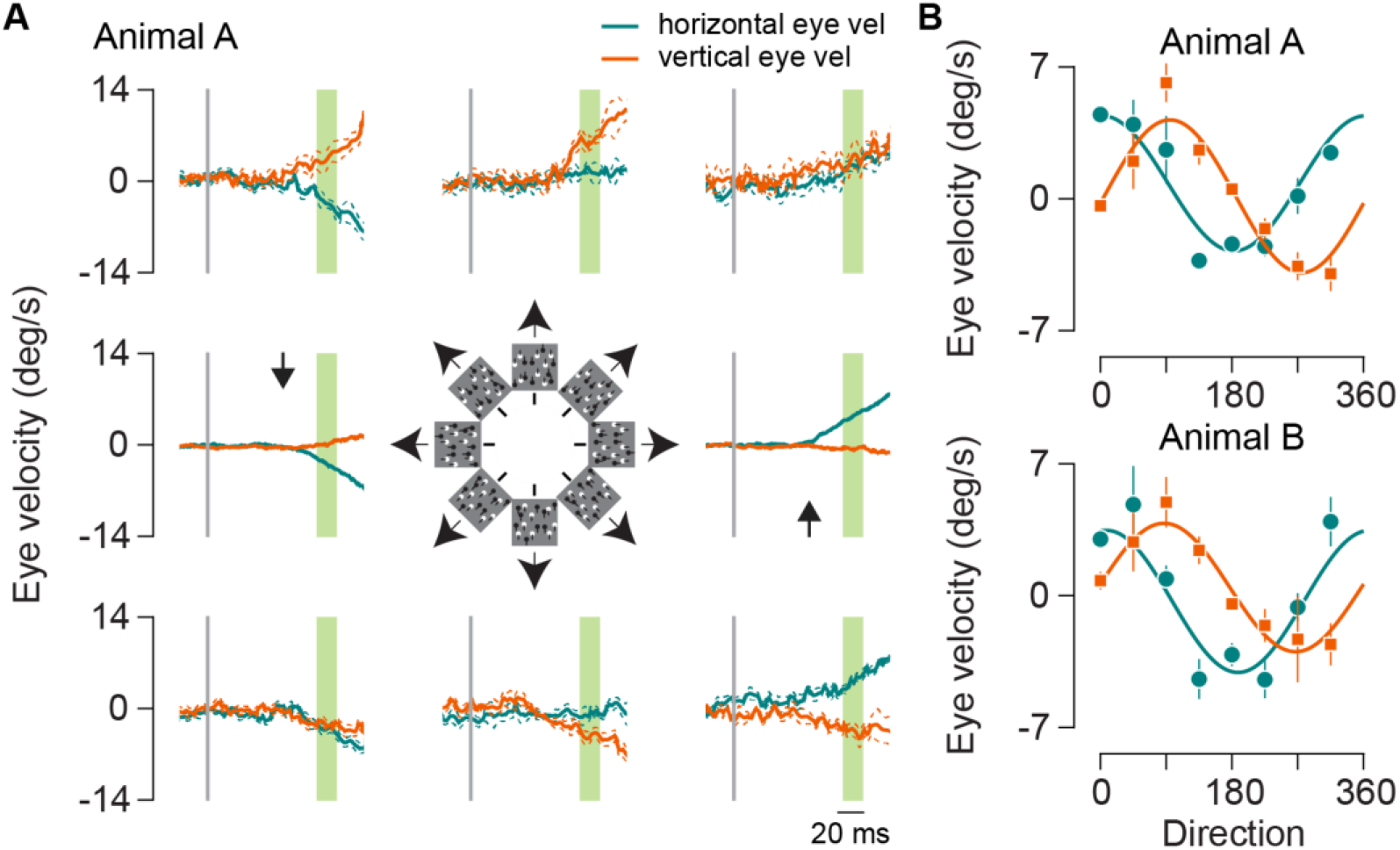
OFR to different directions. A) Mean eye velocities for 8 different dot field directions for animal A. Panels are arranged in counterclockwise order with the right center panel corresponding to rightwards or 0 degrees motion. The cartoon in the center indicates the stimulus motion order. Thick traces in orange and green indicate the vertical and horizontal median eye velocities. Thin traces indicate the s.e.m. around the velocity. Vertical gray line shows the start of motion stimulus. Green highlighted region is used for calculating the amplitude of OFR. Latency was calculated as the mean of the rightwards and leftwards motion condition and is shown by the black arrows. B) OFR amplitude for horizontal and vertical eye velocity for different stimulus motion directions. Upper panel is for OFR in animal A, lower panel is for OFR in animal B. Orange points indicate vertical eye velocity and green indicates horizontal eye velocity. Responses are fitted with sinusoidal function (see methods). Error bars indicate 1 s.e.m.

Having established that marmosets can exhibit ocular following eye movements in response to large field motion, we then examined the range of dot speeds over which OFRs are generated. Using the dot stimulus moving in either the right or left direction we systematically adjusted the speed of the stimuli. The initial eye velocity of the OFRs increased with dot speed, peaking around 25-50 deg/s. Speed greater than that was not as effective in driving ocular following responses (Fig. 3A). The OFR amplitude dependence on speed of motion could be fit with a log Gaussian function (see methods), with comparable mean and standard deviation for the two animals (Fig. 3B, Rightward motion: Animal A: mean = 28.3 deg/s, standard deviation = 2.9 deg/s, Animal B: mean = 41.4 deg/s, standard deviation = 4 deg/s. Leftward motion: Animal A: mean = 33.2 deg/s, standard deviation = 2.6 deg/s, Animal B: mean = 33.7 deg/s, standard deviation = 3.2 deg/s).

**Figure 3:**
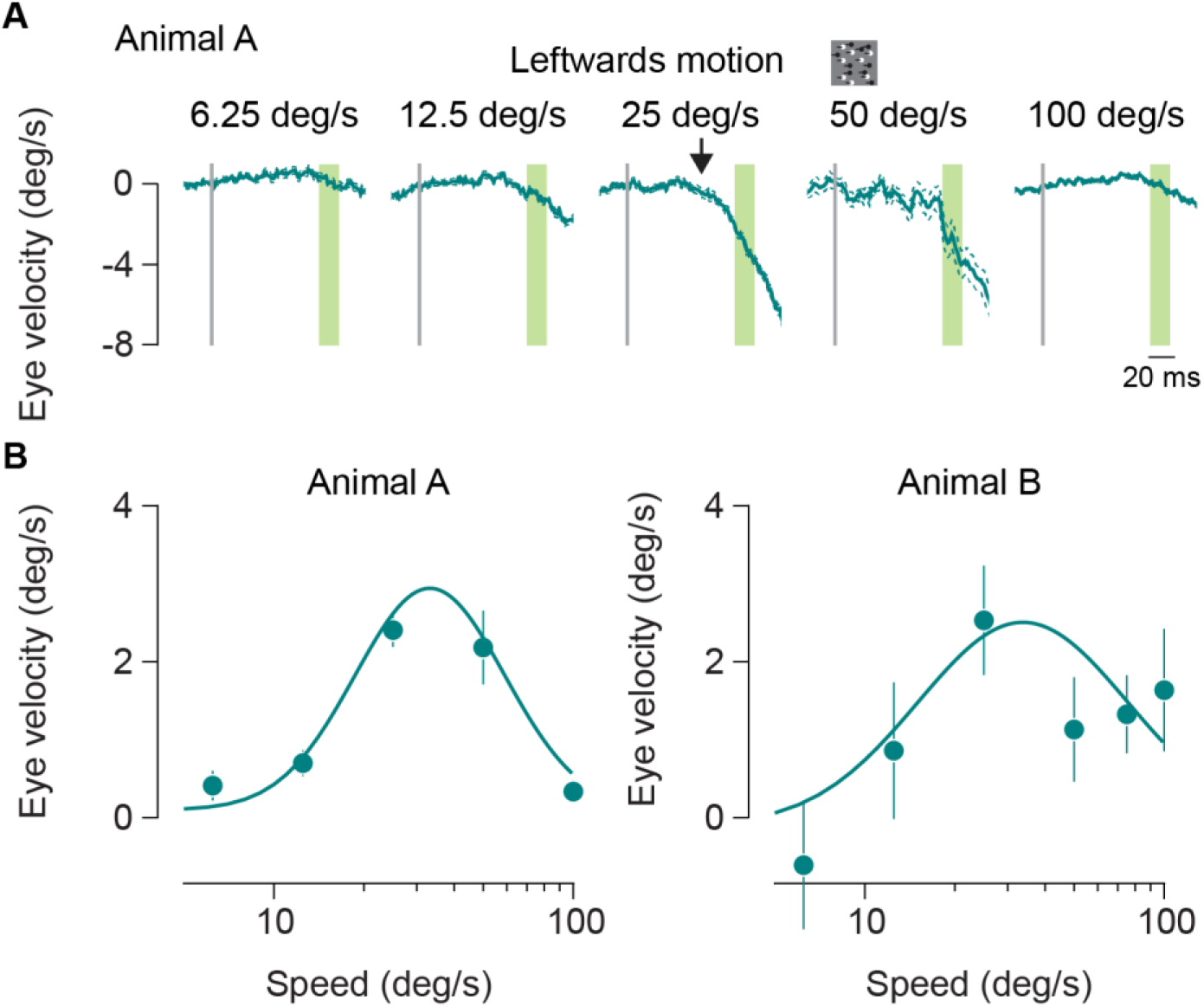
OFR to different speeds. A) Median horizontal eye velocity for dot fields moving leftwards at different speeds. Speed is indicated at top of each panel. Thick line indicates the median eye velocity and thin lines indicate the s.e.m. Gray vertical line shows the start of the motion stimulus. Latency is calculated for the 25 deg/s stimulus, shown by the black arrow. OFR amplitude is average from the period highlighted in green. B) Median OFR amplitude as a function of stimulus speed. Left panel shows responses from animal A and right panel from animal B. Error bars indicate 1 s.e.m. Amplitudes are fit with a log Gaussian function.

### Ocular following responses to single grating motion

Our goal is to examine ocular following responses to combinations of moving gratings. To ascertain that we are using grating parameters that can evoke robust OFRs, we first measured the spatial frequency sensitivity of marmoset OFRs by presenting gratings moving at 25 deg/s at different spatial frequencies. We observe a clear dependence of OFR amplitudes on the spatial frequency of the gratings in both animals. The responses peaked around 0.2-0.5 cycles per degree and declined for both lower and higher spatial frequencies (Fig. 4). The responses were fitted with a log Gaussian function and had comparable mean and standard deviation for different motion directions and across animals. (Rightward motion: Animal A: mean = 0.5 cpd, standard deviation = 3.9, Animal C: mean = 0.3 cpd, standard deviation = 3.2, Animal D: mean = 0.4 cpd, standard deviation = 3.4. Leftward motion: Animal A: mean = 0.5 cpd, standard deviation = 2.9, Animal C: mean = 0.3 cpd, standard deviation = 2.3, Animal D: mean = 0.4 cpd, standard deviation = 2.9). Based on these responses, we have chosen the spatial frequency of 0.4 cpd for the experiments with the plaid stimuli described next.

**Figure 4:**
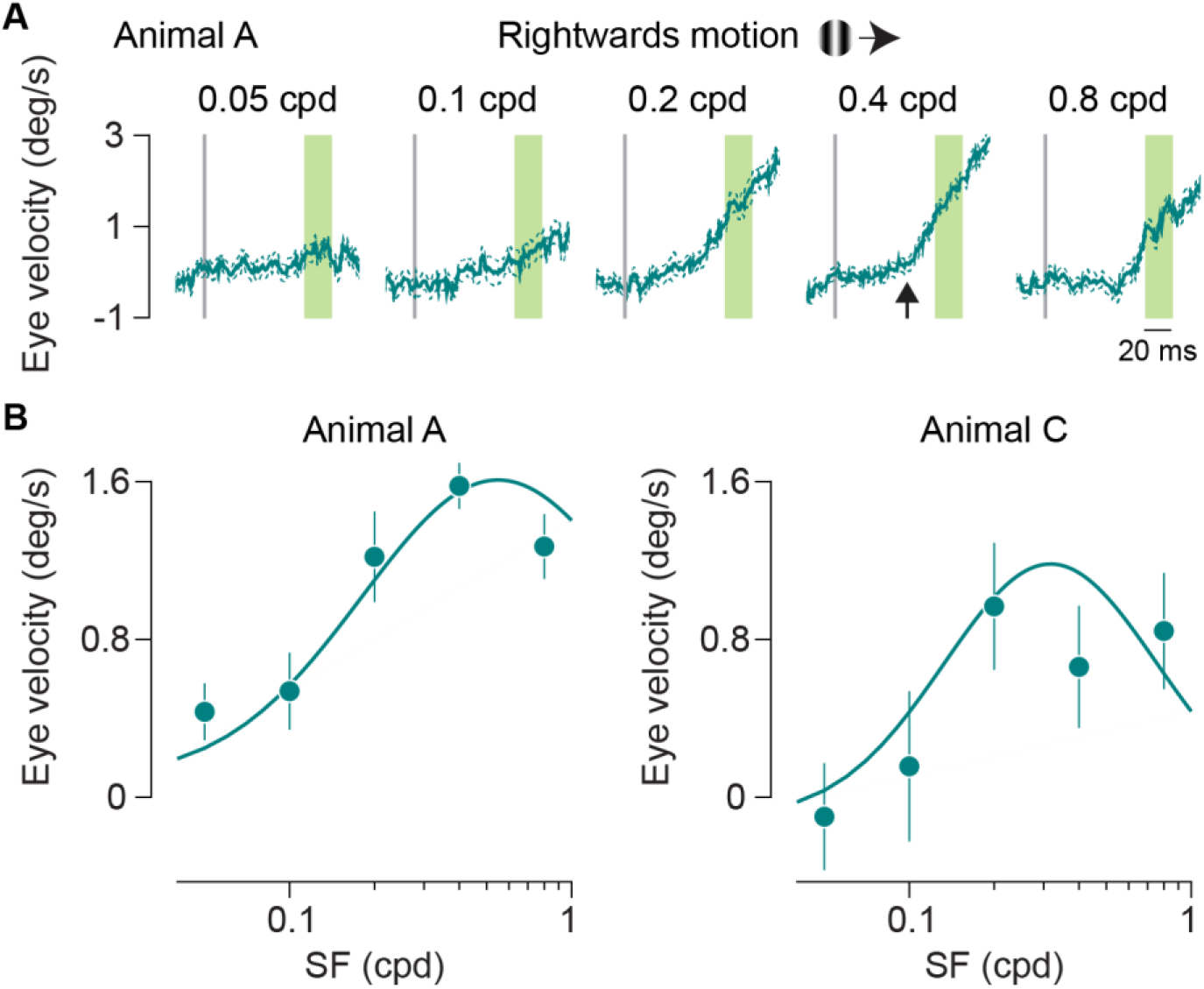
OFR to different spatial frequencies. A) Median horizontal eye velocity for single grating of different spatial frequencies moving rightwards at 25 deg/s. Spatial frequency is indicated at top of each panel. Thick line indicates the median eye velocity and thin lines indicate the s.e.m. Gray vertical line shows the start of the motion stimulus. Latency is calculated for the 0.4 cpd stimulus, shown by the black arrow. OFR amplitude is average from the period highlighted in green. B) Median OFR amplitude as a function of stimulus spatial frequency. Left panel shows responses from animal A and right panel from animal C. Error bars indicate 1 s.e.m. Amplitudes are fit with a log Gaussian function.

### Ocular following responses to plaid motion

Both the dot and grating motion stimuli are composed of strong motion signals consistent with motion in a single direction. Our question, however, revolves around how multiple motion signals are integrated to generate ocular following responses. One approach to assess this is to present two overlapping gratings with distinct orientation and measure how the motion of these grating components is integrated to elicit a behavior such as the OFR (Masson & Castet, 2002). As described earlier, multiple models exist to explain motion integration. Two of the models are vector sum (VS) and vector average (VA), in which the individual motion components are either summed or averaged to give the global motion interpretation. A third model is the intersection of constraints (IOC) model in which each motion component provides a set of constraints to the global motion and the overall motion satisfies all constraints. We use a type II plaid stimulus to test for motion integration in marmosets because it distinguishes between IOC and VA models. This stimulus is formed by superimposing two gratings moving at different speeds along nearby directions. The IOC motion direction in this case falls outside of the two component motion directions (unlike type I plaids). If ocular following eye movement tracks this direction, it offers direct evidence of IOC model of motion integration.

We used a variant of type II plaids, called unikinetic plaids, in which one grating component was stationary, oriented diagonally at 45 degrees, and the other moved at 25 deg/s along the vertical direction, either upwards or downwards. The vector average and vector sum predictions are that motion signals are in the direction of the moving grating component as the second grating is static, whereas the IOC model predicts a computation of motion along an axis of orientation of the stationary grating. Presenting these type II plaids and examining eye movement along the horizontal axis (Masson & Castet, 2002), we can not only test for existence of motion integration but also distinguish between the models used for motion integration computation guiding eye movements. If IOC computations are used to generate OFRs, there should be a smooth eye movement in horizontal direction, whereas if a VA or VS computations are used there should be only vertical smooth eye movements.

We measured smooth eye movements in response to unikinetic plaids and found a component of eye velocity trajectories consistently in the horizontal direction for both marmosets (Fig. 5A), consistent with the IOC model. To estimate whether the horizontal eye velocity observed in response to unikinetic plaids was different from noise, we compared its magnitude to the magnitude of horizontal eye velocity observed in response to single gratings moving in vertical direction. The horizontal component in response to single grating motion along vertical direction gives us an estimate of the noise in the horizontal eye velocity magnitude. We find that the horizontal eye velocity in response to plaid motion was higher than the noise estimate, for both upwards and downwards grating motion in both animals (Fig. 5B, upwards motion t-test animal A, p = 0.04, animal C, p = 0.002, downwards motion animal A, p = 0.1, animal C, p = 0.002), indicating a significant response along the horizontal direction consistent with the IOC model.

**Figure 5:**
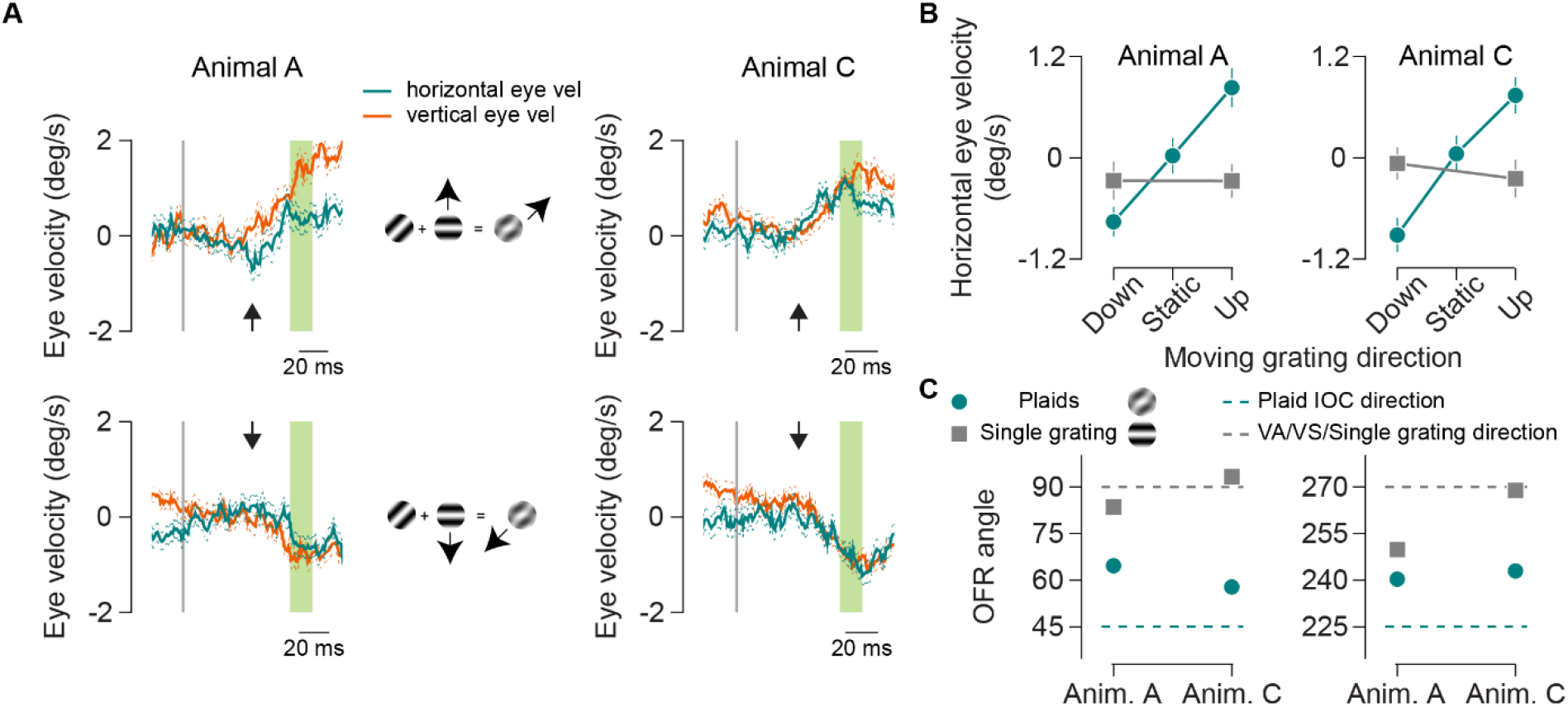
OFR to unikinetic plaids. A) Median eye velocity traces in response to unikinetic plaids motion. Top row shows responses for plaid composed of static diagonal grating and upwards moving grating. Bottom row shows responses for static diagonal and downwards moving grating. First column is for animal A, middle column shows the schematic for plaid composition and third column is for animal C. Thick lines indicate median eye velocities, and thin dashed lines indicate s.e.m. Orange lines represent vertical eye velocity and green lines indicate horizontal eye velocity. Vertical gray lines indicate start of motion. Eye velocity is averaged from the green highlighted section. Latency of responses is indicated by the black arrows. B) Median horizontal eye velocity for plaids along different directions. Left panel is for animal A and right panel is for animal B. Green circles indicate velocity in response to plaids. Gray boxes indicate horizontal velocity for single gratings moving upwards or downwards. Error bars are 1 s.e.m. C) Ocular following angle for plaids and grating stimuli. Left panel is for plaid or grating with moving component along upwards direction and right panel is for plaid or grating with moving component along downwards direction. Green circles are OFR angles for plaid motion and gray boxes are OFR angles are for single grating motion. The direction of the moving grating component is indicated by the dashed gray line and the IOC direction for plaid motion is shown by the green line. Error bars (smaller than points) indicate 1 angular s.d.

At the end of the open loop interval the mean direction of the smooth eye movements was along the diagonal for both animals (Fig. 5C, Plaid with upwards moving grating: circular mean tracking angle for animal A: 64.7 +/- 1.1 angular s.d., animal C: 57.8 +/- 1 angular s.d.. Plaid with downwards moving grating: circular mean tracking angle for animal A: 240.4 +/- 1 angular s.d., animal C: 243 +/- 1 angular s.d.). Note however, that while these horizontal eye movements are consistent with a contribution of the IOC model to the integration of motion trajectories, these angles systematically underestimate the expected angle given the IOC model (45 degrees for plaid with upwards moving grating and 225 degrees for plaid with downwards moving grating).

## Discussion

The new world primate species of marmosets offers exciting advantages as a system to study the link between sensory representations and action: these primates exhibit complex cognitive skills, have lissencephalic brains providing easy access to the neocortex and have well-characterized sensory and motor areas. We have identified the characteristics of the visual motion signals required to elicit ocular following responses in marmosets, which may be used to study the link between neural responses and behavior. In addition, we have demonstrated that marmosets resolve complex motion patterns consistent with an intersection-of-constraints model, as in other primates. Overall marmoset ocular following responses are similar to human and macaques, with species specific differences in peak sensitivities.

Ocular following responses are eye tracking movements generated in response to sudden motion by large stimuli at very short latencies. In humans and macaques, the latency of these responses is reported to be less than 85 and less than 60 ms respectively (Busettini et al., 1994; Gellman et al., 1990; Miles, 1998; Miles et al., 1986). We find that the marmosets also demonstrate short latency tracking movements in response to large field motion with comparable latencies. Within the range of motion parameters we tested, we find that for single grating motions, the maximum response amplitude for marmoset ocular following responses is around 1-2 deg/s. The response amplitude is stronger for stimuli composed of more complex patterns than gratings, such as a field of dots presenting multiple spatial frequencies. In this case, the maximum response amplitudes we observe is around 3-5 deg/s. The lower amplitude with single gratings must therefore reflect poorer activation of involved neurons as the stimulus is restricted to single frequency channel.

We find the marmoset and macaque ocular following spatial frequency selectivity to be similar when compared using gratings. Both species show higher gains at a SF of around 0.3 cpd (macaque results from (Miles et al., 1986)). Area MT, where neurons are selective for motion, is thought to contribute the major sensory drive for ocular following responses in primates (Kawano et al., 1994; Takemura et al., 2007). Unlike the preferred spatial frequency of OFR, the preferred spatial frequency of motion selective area MT neurons of marmosets is lower than that of the macaques (0.2 cpd in marmosets (Lui et al., 2007), 0.6 cpd in macaques (Priebe et al., 2003)). One explanation for the higher SF preference in marmoset OFR could be the higher cone density in marmoset peripheral retina compared to macaque and human retina (Grünert & Martin, 2020; Mitchell & Leopold, 2015; Wilder et al., 1996).

The median optimal spatial frequency for area MT neuron responses in the marmoset is around 0.2 cpd and the median optimal temporal frequency is ~3 Hz (Lui et al., 2007). This sensitivity matches the observed spatial frequency sensitivity of 0.2-0.4 cpd in the ocular following responses. Based on the MT neuron responses, the optimal speed should be around 15 deg/s and we observed there to be a response saturation at speeds beyond 25 deg/s. Given the overlap in the peak sensitivity for these motion parameters between marmoset MT neuron responses and marmoset MT ocular following responses, it is possible for these MT responses to be underlying the ocular following responses in the marmoset. The small discrepancies in peak sensitivities between OFR and neural responses may be due to contribution of MST neurons, which in macaques is known to have different sensitivity than MT (Miura et al., 2014). Finally, the gain in ocular following response to motion of different speeds, saturates earlier in the marmoset than the macaque. This may be a result of smaller head sizes and higher prevalence of head movements (not possible under head fixed conditions) in marmosets when they are tracking motion (Pandey et al., 2020).

Human and macaque observers have been shown to successfully track the integrated trajectory of spatial patterns formed of multiple motion components (Barthélemy et al., 2008; Masson & Castet, 2002). Marmosets also track such integrated motion when presented with plaid patterns. In the human and macaque studies, it was observed that the initial tracking component would be in response to the local motion component whereas the response to the integrated motion has a slightly longer latency and tends to follow with a delay of 20 ms. In our records, the distinction between tracking directions in the early time intervals has been difficult to assess as the initial marmoset eye movements are small, causing the initial tracking direction estimates to be noisy.

The neural basis underlying the responses in the integrated motion direction are less clear. Marmoset MT, just like macaque MT, is composed of pattern cells, component cells, and cells in between the spectrum (Solomon et al., 2011). It remains unclear as to how this diverse MT population signals the visual motion direction as a whole (Quaia et al., 2022). Previous studies have demonstrated contribution of multiple motion sensing mechanisms, such as based on first order motion, second order motion as well as features (Barthélemy et al., 2008; Lorenceau et al., 1993; Masson et al., 2000). Motion detection/integration can occur differently in each of these channels and together may contribute to the ocular following eye movements. It is also likely that selectivity for integrated global motion is enhanced in circuits following area MT which finally contribute to guiding ocular following direction. This does not negate the possibility for subcortical contribution to ocular following responses (Inoue et al., 2000). The ease of access to cortical and circuitry in marmosets, the ability to train them to perform simple eye movement dependent tasks and the similarity of visual pathway with macaques and humans, makes them an ideal model to study the neural basis of motion computations and explore how they guide behavior such as eye movements.

## Acknowledgements

We would like to thank Guillaume Masson for advice on setting up the ocular following task and to Samuel Poelker-Wells and Allison Laudano for assistance with animal care.

